# Onset of Mismatch Repair by the Human Mismatch Repair Protein, MutSbeta for a Biosensing Device

**DOI:** 10.1101/2024.01.31.578329

**Authors:** Jack Devlin, Jenna Madigan, Isaac Macwan

## Abstract

Deficiency of MutSbeta protein has been proven to be a cause of Lynch Syndrome, which leads to hereditary colorectal cancer. In the absence of MutSbeta, or the energy source to initiate the conformational change of MutSbeta, the DNA repair pathway would not detect the DNA base pairing errors resulting in a failure to proceed. To examine the essential role of MutSbeta and the interface between MutSbeta and a mismatched DNA strand, 500ns of molecular dynamics simulations were performed and the results were compared with the controls. It is found that during the first 100ns, the outermost domain of MutSbeta (chain BP4) gets hold of the DNA, and the inner domains (AP1 and BP1) prepare to scan the DNA strand. An RMSD of 5-7Å for the control MutSbeta compared to 3-6.5Å in the presence of a mismatched DNA indicate that starting ∼75ns, MutSbeta stabilizes and initiates the scanning of the mismatched DNA. The reduction of the distance between the two biomolecules by ∼16Å, increase of Van der Waals energies by ∼75 kCal/mol and a crucial role played by the interfacial water molecules and their hydrogen bonds during the next 100ns also supports the manipulative nature and initiation of scanning by MutSbeta.

**Author Summary:** About 1 and 23 men and 1 and 26 women have the chances of developing colorectal cancer and the chances only increase with old age. Cancer typically develops from mutated DNA, which emphasizes the importance of understanding how proteins a part of the DNA repair pathway process interact with a mutated strand of DNA. One such protein, MutSbeta, is the first protein that initiates the repair pathway process and scans DNA for any mutations. As the initiator, a deficiency of MutSbeta leads to higher chances of mutated DNA going undetected, and therefore forming cancerous cells. Using 3D simulations, we analyzed the interactions between MutSbeta and a mutated DNA strand. We found that the two outer domains, AP4 and BP4 grab hold of the DNA strand while the inner clamps, AP1 and BP1 scan the DNA strand. It was also discovered that Van der Waals play a more dominant role compared to electrostatics in terms of the driving force behind the interactions between DNA and MutSbeta. Overall, our data supported the attractive nature of MutSbeta towards DNA and its stability as it initiates the scanning process. Furthermore, this data was used to verify the presence of MutSbeta for a biosensor that was built previously; owing to the significance and applicatory effects of this work.

## 1. Introduction

With an increased interest in cellular disease and cancers, the study of DNA becomes essential to consider due to its complexity and the process of its replication, where errors related to the mismatch are prone to happen. Many internal and external factors can cause DNA damage, whether it is errors in replication or external factors such as from exposure to radiation, other cellular materials, or chemical exposure [1]. These errors then compound as strands multiply quickly and cellular mechanisms must counteract via DNA repair to prevent further damage. It is known that the process of DNA replication occurs through a complex mechanism at the cellular level [2]. Methods of DNA repair include nucleotide excision repair, base excision repair, and the mismatch repair pathway [3]. In a mismatch, the errors result from an improper replication by DNA polymerase and show up in the form of substitution of incorrect bases and insertion and deletion of bases [4]. Mutation rates are noticeably elevated when a defective MMR pathway is found [5]. With proper functioning of the MMR pathway, these errors can be repaired and will keep the original, intended makeup of the strand [6]. The replication process in humans is remarkably accurate but can occur randomly. The proteins that are essential for this entire process include MutS, MutL, an exonuclease, DNA polymerase, and DNA ligase [7].

The protein of interest MutS is composed of two different heterodimers: MutSalpha (MSH2-MSH6 heterodimer) and MutSbeta (MSH2-MSH3 heterodimer) [8]. This study deals with the role of MutSbeta (MSH2-MSH3) component. The structure of MutSbeta involve twelve chains AP1-AP6 and BP1-BP6. AP1 and BP1 chains make up the inner clamp and the AP4 and BP4 chains make up the outer clamp [8]. The outer clamp grabs the free end of the DNA and serve as the docking spot for the protein’s inner clamps to scan the DNA. These two clamps were the focus of study as their interaction with the mismatched DNA continued. The modeled DNA that we studied had 50-51 base pairs on either strand where resid84 was removed to model the mismatch [8]. MutL forms a complex with MutS after the mismatch is identified by the heterodimer and helps to hold the MutS at the site [9].

Human MutsB slides along behind the DNA polymerase on the newly synthesized DNA to recognize any errors made in the smaller insertion loops (1-4 base pairs) of the mismatch repair system [10]. According to Modrich, a nick on the strand must be present to recruit the MutS complex to the site of a mismatch [11]. This signal is not required to be immediately next to the errors but in its vicinity to signal a need for the pathway to ensue. If the MutsB recognizes an incorrect base pairing or sequence, MutL will then be recruited to the scene to mark the area to be removed and the exonuclease will follow to remove the unwanted portion [8]. The process continues with DNA polymerase filling in the missing amino acids and DNA ligase sealing the new sequence to the original. It is important to note that the length of the replaced DNA is usually about 1000 bases in length, which differentiates MMR from the processes of Base Excision Repair, which is only 1 nucleotide replacement [12], and Nucleotide Excision Repair, which provides about 24-30 new nucleotides [13]. MutsB acts on an energy source known as adenosine diphosphate, or ADP, to initiate the repair pathway process; without ADP, the necessary conformational change of MutsB will not take place [14]. Lack of this energy would not allow the binding of MutS and would inhibit the rest of the mismatch repair sequence; inactivation of this protein increases the chances of mutations to occur [15]. The sliding clamp setup forms when ADP binds and the clamp then scans to find the error [16]. There is little knowledge on how these clamps of MutSbeta bind and interact with a mismatched DNA strand. In the past, researchers have hypothesized that insertion deletion loops in the DNA will be distorted in the backbone of the sequence [17]. It is also expected that a bend or loop in the DNA will be oriented at an angle of 45 to 60 degrees [18].

When the MutSbeta component is absent, mutations will multiply rapidly. The result of this uncontrolled and mutated cell growth is cancer, specifically Lynch Syndrome [5]. Hereditary Nonpolyposis Colorectal cancer (HNCC) results from Lynch Syndrome and the mutation or incorrect function of MutSbeta [19]. With this background, this study focuses on the mismatch occurring due to the incorrect substitution of DNA bases, specifically base T on a DNA backbone and the trajectory that the mismatch repair protein MutSbeta will take to initiate scanning of the mismatched DNA. Biosensors using graphene have been studied previously [20,21,22], but none to quantify the presence of MutSbeta to address the formation of HNCC through the initial DNA repair pathway steps. With this being said, from a biosensing perspective, having a platform that captures the presence of MutSbeta will drastically improve the understanding of Lynch Syndrome.

From a previous study, a biosensor platform with graphene and avidin docks one end of a biotinylated mismatched DNA through the avidin biotin chemistry with the mismatch engineered at the free end of the DNA strand [23].Keeping this setup in mind, which allows access for the MutSbeta to latch and find the mismatch on the strand, the present study models the mismatched DNA to freely interact with the MutSbeta at one end while its other end is fixed. From the analysis of the simulated trajectories for the controls and overall system, the onset of DNA repair is anticipated through changes in the secondary structure of the clamps, a reduced distance between the two molecules’ center of mass and from other data including stability and energetics such as RMSD (Root Mean Square Deviation), hydrogen bonds, salt bridges, and interaction energies involving conformational, electrostatics and Van der Waals. Role of interfacial water molecules and their hydrogen bonds is also highlighted. Where a goal of this study is to determine which forces play a major role in the interaction between the mismatched DNA strand and MutSbeta in order to support the biosensing system.

## 2. Materials and Methods

The interactions between human MutsB and a mismatched DNA strand were observed at the molecular level using a molecular graphics tool called Visual Molecular Dynamics (VMD) [24]. A molecular simulation program called NAMD (Nanoscale Molecular Dynamics) is used to simulate the interactions between a human mismatch repair protein, MutSbeta (PDB 3THY) that is complexed with an IDL (imidazo[1,2-a]pyridin-3-ylacetic acid) of 2 bases (loop 2) and ADP. A 50-51 base pair strand of DNA was modeled using an online DNA modeler tool called: The Sequence Manipulation Suite [26], and VMD was used to create the mismatched base pair by editing the PDB file, specifically the mismatch site, the Resid84 on the mismatched DNA. One end of the DNA strand was also fixed to mimic a biosensor surface having immobilized mismatched DNA as a probe. A control group was first simulated for 100ns each for the mismatched DNA and MutSbeta to quantify their stability individually. The two molecules were then combined into a single overall system in a water box with neutralizing concentration of NaCl ions to study their stability in the presence of each other and energetics as well as role of interfacial water molecules.

A water box containing 380,750 water molecules was created with a size of 115×115x329 Å using TIP3P [26] water model. All-atom simulations were carried out for 500ns using NAMD [27] and CHARMM [18] force field on an Intel Core i9 cluster with a total of 36 cores, and NVIDIA GeForce RTX 2080 GPU. The boundary conditions were assumed at a constant temperature of 310K through Langevin thermostat and a pressure of 1atm by Langevin barostat [28]. The minimization and equilibration of the system was performed for 5000 steps and 500,000 steps respectively with a timestep of 2fs for all simulations. To analyze the stability of the system, RMSD, salt bridges, and center of mass were analyzed using the built-in plug-ins and TCL scripts within VMD. The Van-der-Waals interactions are evaluated in terms of Lennard-Jones potential that typically describe the ex-change-repulsion and the dispersion attraction of two molecules [29]. Coulomb potential was used for the evaluation of electrostatics, non-bonding energies via Ewald Summation (also known as PME) [30]. All data is plotted using Origin Pro [31].

## 3. Results & Discussion

Essential components of the relationship between MutSbeta and mismatched DNA are seen through the analysis of stability and energetics.

In Figure 1, the schematic shows the visual representation of both DNA and MutSbeta. In the first image at 0 ns, the two components begin within ∼15 Å from each other to promote a quick interaction between the two. Based on the simulated trajectory, it turned out that several hundred nanoseconds worth of data was needed to begin to examine changes in the system’s energy and configuration. The fixed nature of the left end of the mismatched DNA promoted a solid docking ground on the loose end for the MutSbeta to latch. As the progression in Figure 1 continues downward, the leftward arrow show the progression of MutSbeta down the DNA strand towards the mismatch. As mentioned in the figure, the outer clamps AP4 and BP4 complete the initial grab of DNA and help the protein to begin the docking and scanning process. At the end of 500ns, the DNA and protein are close enough to scan for the mismatch. These figures provide a visual insight into the onset of the mismatch repair process and the numerical data further substantiates these claims. The secondary structure analysis of the four chains AP1, AP4, BP1 and BP4 that make up the inner and outer clamps during the 500ns simulation run can be found in the supplementary figures S1 to S4.

**Figure 1.**
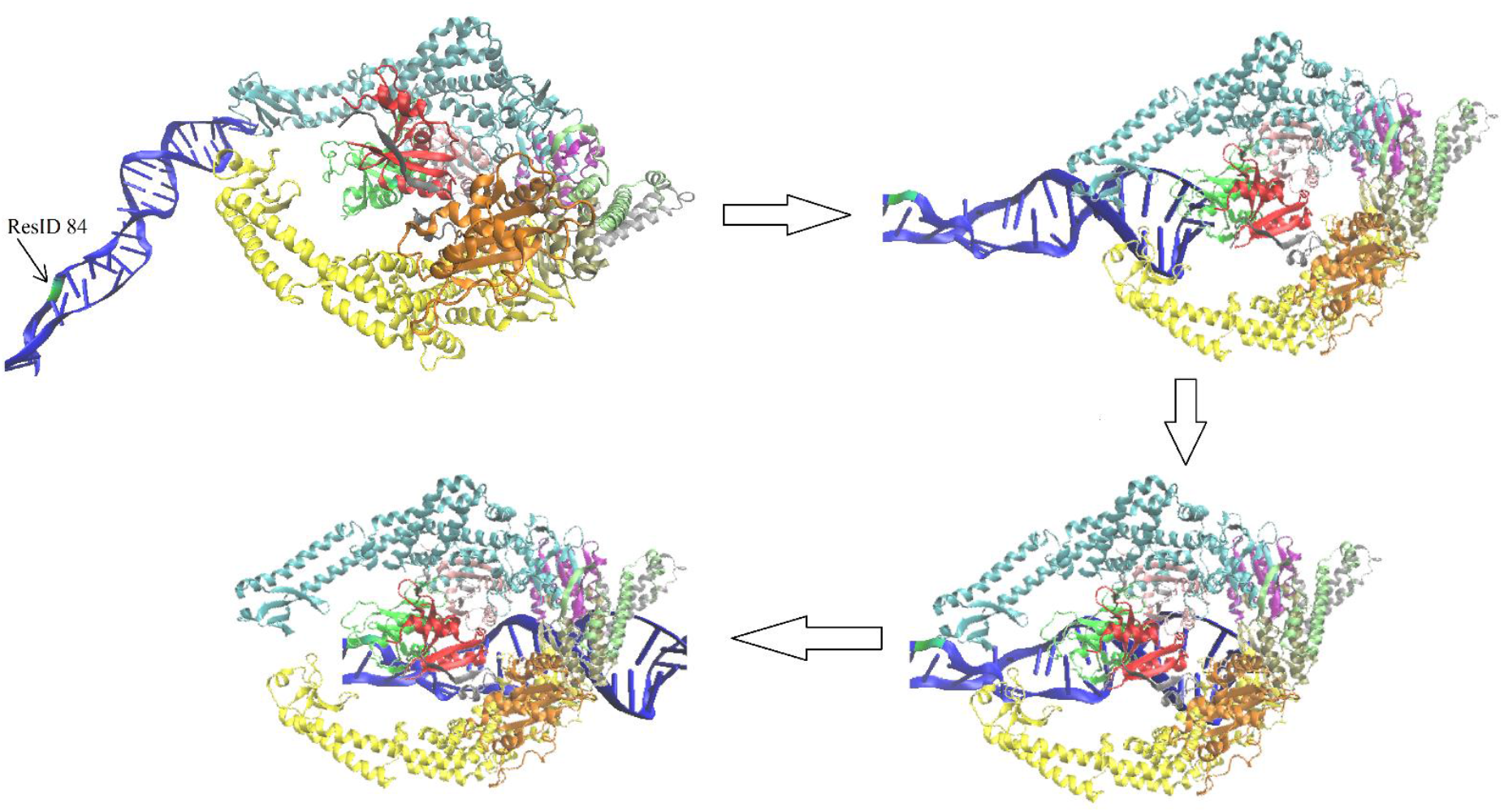
Schematic Depicting the Relationship and Movement of DNA and Protein. At 0 ns (top left), the docked DNA begins to be approached by MutSbeta. At ∼100 ns (top right), MutsBeta gets hold of the mismatched DNA undergoing conformational changes but not yet scanning. At ∼200 ns (bottom right), the scanning process initiates when BP4 (teal) interacts with the mismatch site. From 200 ns onwards, BP4 segment continues to interact with the mismatched DNA as both AP4 and BP4 (yellow and teal) continue towards the mismatch (bottom left).

To continue, the examination of the center of mass provides insight into the potential reduction in distance as the molecules move closer to each other and allow for the DNA repair to proceed correctly. Figure 2A shows that between the centers of mass of each molecule, they begin at ∼158 Å apart and slowly, a reduction in distance occurs down to ∼140 Å at the end of simulation run of 500 ns. This data is consistent with the expected values because the system is made up of about four hundred thousand atoms, meaning the centers of mass should still both be fairly separated when the two are interacting. On the other hand, it is essential to consider the location of the mismatch we aim to examine; this mismatch is near the free end of the DNA. On a deeper level, the center of mass analysis was compared specifically from MutSbeta’s center of mass to the mismatch site, which is the resid84 on the mismatch DNA. This data reinforced the first center of mass analysis as it also showed a substantial distance reduction from ∼135 Å to ∼114 Å. This distinction of about 21 Å shows a more significant drop in the distance between MutSbeta and the mismatched DNA and supports the idea that the onset of interactions is beginning to occur between MutSbeta and mismatched DNA.

**Figure 2:**
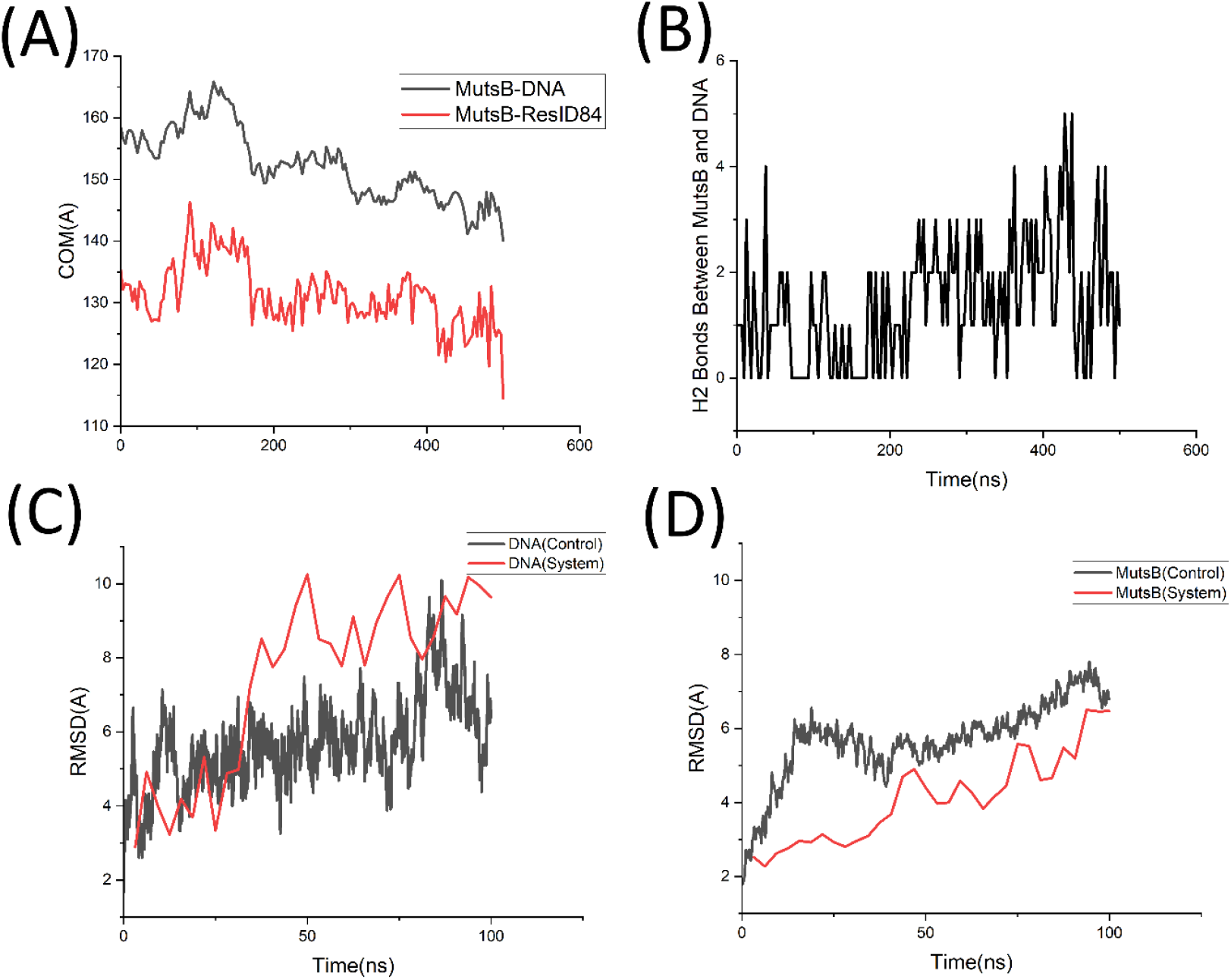
(A) Center of Mass Analysis between MutSbeta & mismatched DNA, and between MutSbeta & mismatch site at Resid84 on the mismatched DNA (B) Number of Hydrogen Bonds between MutSbeta and DNA (C) RMSD of mismatched DNA in the absence (control) and presence (system) of MutSbeta (D) RSMD of MutSbeta in the absence (control) and presence (system) of mismatched DNA.

In Figure 2B, the number of hydrogen bonds allow for the examination of bonds between hydrogen and oxygen atoms at the interface of MutSbeta and the mismatched DNA. The interactions on a molecular level down to the simplest atoms hint at more complex interaction at the level of protein and DNA. As observed from figure 2B, up until 200ns, the number of hydrogen bonds fluctuated between 0 and 2 mostly with a few spikes indicating very short lifetimes of the bonds. However, as time progressed, the number of bonds jumped to rest between 0-5 with longer lifetimes of ∼3 bonds on an average between MutSbeta and the mismatched DNA. This uptick in bonding helps explain the closer distance between atoms and their ability to work together in finding a mismatch and therefore signaling for its scanning and eventual repair.

Furthermore, Figures 2C and 2D highlight the RMSD to quantify the stability of the MutSbeta and the mismatched DNA. As observed in Figure 2C, the RMSD of mismatched DNA control hovers around the value of ∼5 Å, whereas that of the mismatched DNA in the presence of MutSbeta stabilizes around ∼9 Å, which represents a larger deviation owing to its conformational changes in the presence of MutSbeta. It is to be noted that the free end of the DNA may also cause some oddly high RMSD values. In figure 2D, a similar RMSD analysis of MutSbeta in the presence and absence of mismatched DNA is performed where the data highlights the fact that the RMSD of the control group deviates much more compared to the system. This comparison further suggests that the protein settles down as it latches onto the target DNA as seen from its RMSD tapering off at ∼6 Å after the initial spikes by the end of the first 100ns. RMSD data for MutSbeta and mismatched DNA for the entire 500ns is provided in the supplementary figure S5, which further points out that at ∼200ns, when the protein starts scanning the DNA, its RMSD dips to ∼4 Å and stabilizes there.

All in all, these stability interactions provide solid evidence for the initial interactions between the mismatched DNA and MutSbeta. As the analysis proceeds, the case to support these interactions becomes more obvious with interaction and conformational energies, analysis of optimal distance between MutSbeta and mismatched DNA, and role of interfacial water molecules with respect to the polar residues of the MutSbeta.

Figure 3 presents data related to the energies of the mismatched DNA, MutSbeta, and a combination of the two for proper examination of their influence on each other in the system. These interactions, while weak (being nonbonding), have a significant impact on the final orientation and conformation of both, the amino and nucleic acids for MutSbeta to get in a proper position to initiate the scanning of the mismatched DNA. In Figure 3A, the nonbonding energy is overlaid for the MutSbeta for both in the absence and presence of the mismatched DNA. It is observed that initially the nonbonding energy of the MutSbeta in the absence of mismatched DNA stays around ∼41,750 kcal/mol whereas in case of the presence of mismatched DNA, the nonbonding energy of the MutSbeta, though similar to the control case initially, dips starting ∼35ns and continues to decrease to ∼43,250 kcal/mol by the end of the first 100ns. This relatively larger value for nonbonding energy indicates the presence of attractive forces within MutSbeta as it approaches the mismatched DNA. The nonbonding energies for MutSbeta in the presence of mismatched DNA for the entire 500 ns is provided in supplementary figure S6. Figure 3B shows the Van Der Waals interactions between MutSbeta and the mismatched DNA. As observed from Figure 3B, starting ∼150ns, the VDW energy values continue to decrease significantly until ∼220ns where it approaches average value of ∼75 kcal/mol. The negative values indicate the attractive nature of the interface between MutSbeta and mismatched DNA. In Figure 3C, the electrostatic energy within the mismatched DNA in the absence and presence of MutSbeta is shown and as observed, the electrostatic energy fluctuates around ∼9250 kcal/mol for both the cases indicating the not so significant role of electrostatic energy in the mismatch DNA scanning process. Similarly, in Figure 3D, the electrostatic energy of MutSbeta in the absence and presence of mismatched DNA shows that it is around -35000 kcal/mol for both the control and the system simulations. This data indicates that the charge and the distance may be playing some role in the electrostatic energy but overall, it seems that electrostatic energy is not the major driving force behind the interactions between MutSbeta and the mismatched DNA. The electrostatic energies for the MutSbeta and mismatched DNA for the entire 500ns are provided in supplementary figures S7 and S8.

**Figure 3:**
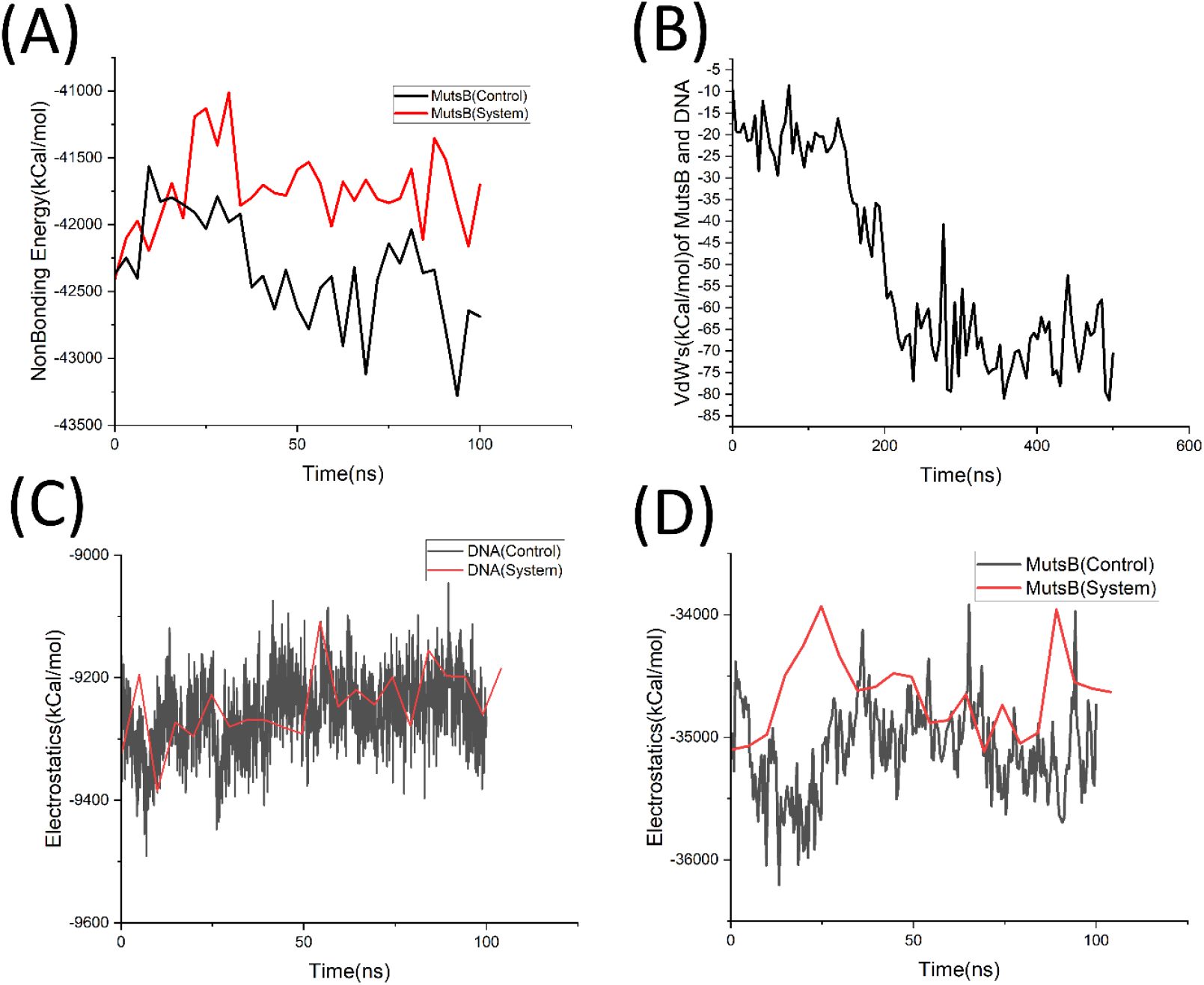
(A) Nonbonding Energy of MutSbeta in the absence and presence of mismatched DNA (B) Van der Waals Energy between MutSbeta and mismatched DNA (C) Electrostatics energy of the mismatched DNA in the absence and presence of MutSbeta (D) Electrostatics energy of MutSbeta in the absence and presence of mismatched DNA.

When comparing the graphs of figure 3 to Figures 4A and 4B it can be observed that Van der Waals, and not electrostatics, plays a key role in the attraction of MutSbeta towards DNA. In figures 4A and 4B, we see that the Van der Waals energy is stable at around -7100 kcal/mol and around -50 kcal/mol for MutSbeta and DNA respectively. The high attraction of Van der Waals for MutSbeta and low attraction of Van der Waals from DNA, combined with the inverse graph of electrostatics of MutSbeta system relative to the control shows that Van der Waals plays a more dominant role in the interaction between the two molecules.

**Figure 4:**
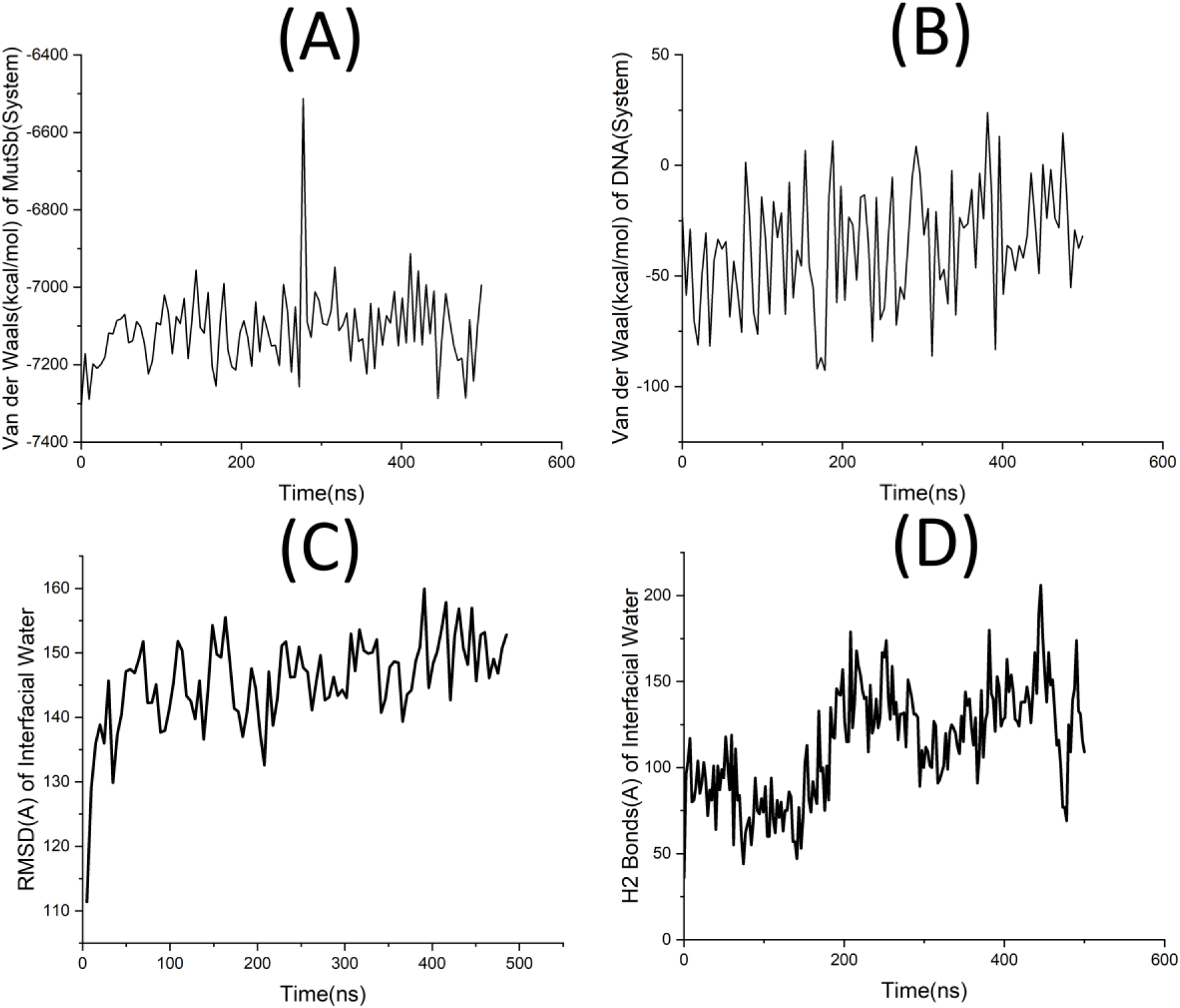
A) Van der Waals of MutSbeta from the system B) Van der Waals of DNA from the system C) RMSD of Interfacial water between MutsB and DNA D) Hydrogen bonds of Interfacial water between MutsB and DNA

After considering these energetic values, it is found that these energies reinforce the stability of DNA and MutsB when in the system as they approach each other. It also adds to the fact that the charges of the two molecules play a huge role in how they interact with each other.

As we continue to analyze interfacial water, the RMSD of this system component is essential to examining more stability in the region of interactions. Here, Figure 4C highlights the RMSD hovering between 135 Å and 155 Å indicating the stability of the interfacial water molecules between MutSbeta and DNA and hence the stability in the interactions of MutSbeta when encountering the mismatched DNA. In Figure 4D, hydrogen bonds between the interfacial water molecules play a crucial role as examined in the area of water between the DNA and protein. In this area shared between DNA and protein, the consideration of water is important because water can have key interactions due to its charge and ability to bond. The number of hydrogen bonds seem to increase slightly from 0-500 nanoseconds. The data begins by possessing about 100 hydrogen bonds and ends up finishing around 125 hydrogen bonds. This reinforcement furthers the ability to confirm increased interaction between MutSbeta and DNA.

The conformational change of MutSbeta was also observed via the salt bridges that were either formed, broken, or sustained throughout the simulation. They are seen in tables 1 and 2 where 18 common salt-bridges were found between the control and system and of these 18, five of them changed conformation. The ones to take note are: GLU16-LYS73, ASP859-ARG854, GLU603-LYS30, GLU877-ARG846, and GLU877-ARG881. These salt-bridge are evidence that MutSbeta undergoes a conformational change to latch onto the DNA strand and is more stable in the presence of DNA compared to its absence. As observed from Table 1 and Table 2, the number of salt bridges which are either formed or broken within the two clamps AP and BP in the absence of DNA are 18 whereas those in the presence of DNA are 12. This reduction in the dynamics of salt bridges indicates that MutSbeta is more stable in the presence of DNA, thereby triggering the molecular events to initiate the mismatch repair.

**Table 1:** Control salt-bridges for AP1, AP4, BP1, and BP4 for MutsB that are either broken, formed, or sustained.

| Salt Bridge Control | Broken | Formed | Sustained |
| --- | --- | --- | --- |
| **AP1** |  |  |  |
| **ASP38-LYS104** |  |  | ✓ |
| **ASP49-LYS29** |  |  | ✓ |
| **GLU16-LYS73** |  |  | ✓ |
| **GLU56-ARG96** |  |  | ✓ |
| GLU86-LYS82 | ✓ |  |  |
| **GLU101-ARG35** |  |  | ✓ |
| **AP4** |  |  |  |
| **ASP386-ARG340** |  |  | ✓ |
| ASP459-ARG482 |  | ✓ |  |
| ASP660-LYS661 |  | ✓ |  |
| ASP706-LYS627 |  | ✓ |  |
| GLU358-ARG355 |  | ✓ |  |
| **GLU364-ARG621** |  |  | ✓ |
| GLU467-LYS567 |  | ✓ |  |
| GLU643-ARG638 |  |  | ✓ |
| GLU658-LYS661 |  |  | ✓ |
| GLU701-ARG631 |  | ✓ |  |
| **BP1** |  |  |  |
| **ASP236-LYS294** | ✓ |  |  |
| **ASP253-LYS231** |  |  | ✓ |
| **GLU302-LYS307** | ✓ |  |  |
| **BP4** |  |  |  |
| **ASP586-ARG839** |  | ✓ |  |
| ASP611-ARG565 |  |  | ✓ |
| ASP657-ARG660 | ✓ |  |  |
| ASP708-LYS704 |  |  | ✓ |
| **ASP793-ARG706** |  |  | ✓ |
| **ASP859-ARG854** | ✓ |  |  |
| **GLU603-LYS830** | ✓ |  |  |
| **GLU613-ARG565** |  |  | ✓ |
| GLU642-LYS608 | ✓ |  |  |
| GLU692-LYS707 | ✓ |  |  |
| GLU742-ARG787 |  | ✓ |  |
| **GLU746-ARG770** | ✓ |  |  |
| GLU845-ARG846 | ✓ |  |  |
| **GLU877-ARG846** |  |  | ✓ |
| **GLU877-ARG881** |  |  | ✓ |

**Table 2:**
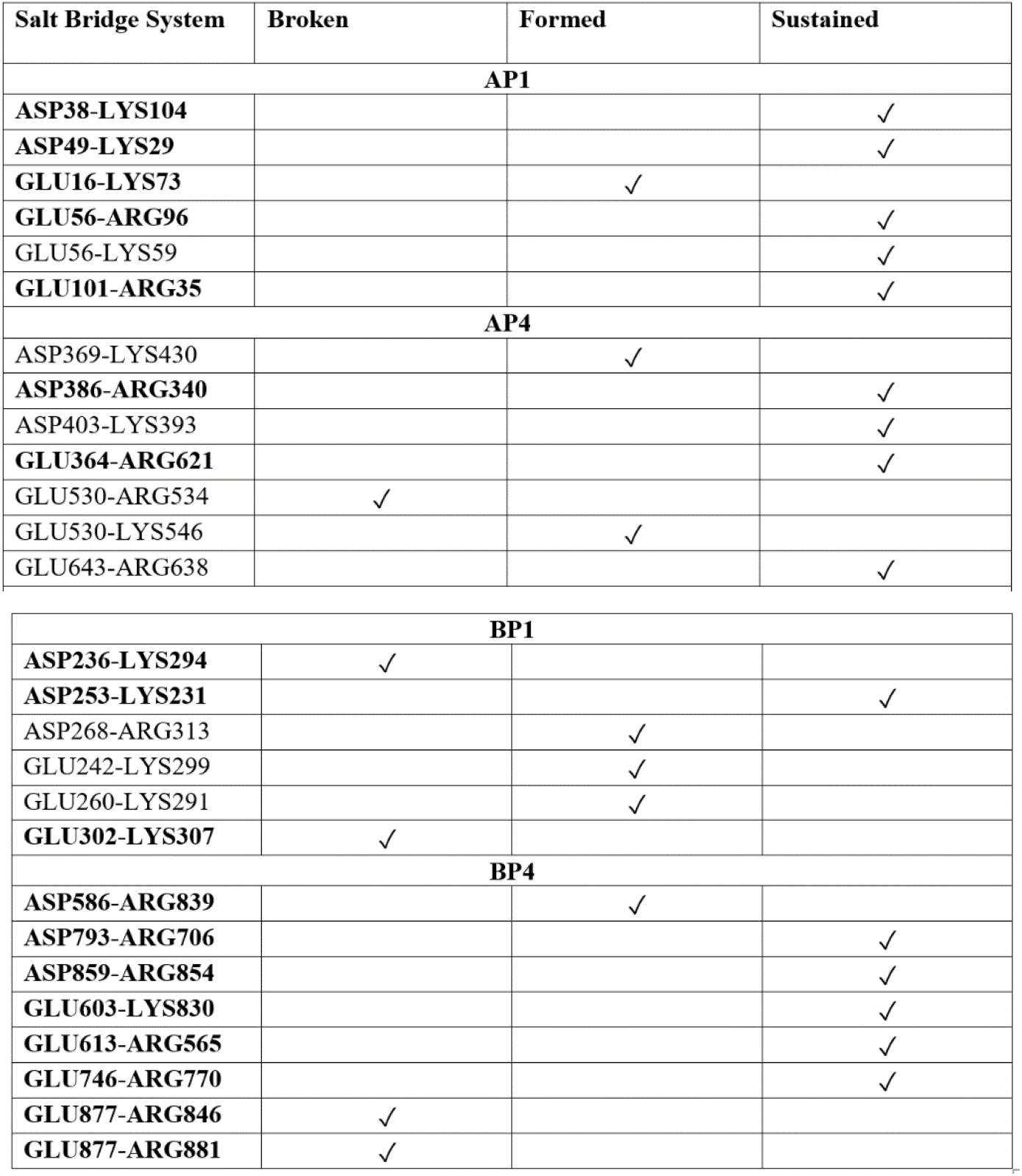
System salt-bridges for AP1, AP4, BP1, and BP4 for MutsB that are either broken, formed, or sustained (**Bold** salt-bridges are ones that appear in both the control and system)

| Salt Bridge System | Broken | Formed | Sustained |
| --- | --- | --- | --- |
| **AP1** |  |  |  |
| **ASP38-LYS104** |  |  | ✓ |
| **ASP49-LYS29** |  |  | ✓ |
| **GLU16-LYS73** |  | ✓ |  |
| **GLU56-ARG96** |  |  | ✓ |
| GLU56-LYS59 |  |  | ✓ |
| **GLU101-ARG35** |  |  | ✓ |
| **AP4** |  |  |  |
| ASP369-LYS430 |  | ✓ |  |
| **ASP386-ARG340** |  |  | ✓ |
| ASP403-LYS393 |  |  | ✓ |
| **GLU364-ARG621** |  |  | ✓ |
| GLU530-ARG534 | ✓ |  |  |
| GLU530-LYS546 |  | ✓ |  |
| GLU643-ARG638 |  |  | ✓ |
| **BP1** |  |  |  |
| **ASP236-LYS294** | ✓ |  |  |
| **ASP253-LYS231** |  |  | ✓ |
| ASP268-ARG313 |  | ✓ |  |
| GLU242-LYS299 |  | ✓ |  |
| GLU260-LYS291 |  | ✓ |  |
| **GLU302-LYS307** | ✓ |  |  |
| **BP4** |  |  |  |
| **ASP586-ARG839** |  | ✓ |  |
| **ASP793-ARG706** |  |  | ✓ |
| **ASP859-ARG854** |  |  | ✓ |
| **GLU603-LYS830** |  |  | ✓ |
| **GLU613-ARG565** |  |  | ✓ |
| **GLU746-ARG770** |  |  | ✓ |
| **GLU877-ARG846** | ✓ |  |  |
| **GLU877-ARG881** | ✓ |  |  |

Diving deeper into the analysis, exhaustive measures were performed to gain the most insight from the data. Interaction energy and the number of atoms is compared in the form of a ratio to determine the optimal distance between the mismatch DNA and MutSbeta for maximum interaction. The different legend points on the graphs in Figure 5 highlight the difference in the number of atoms with and without hydrogen. Due to the small size of hydrogen atoms, their addition to the mix causes the number of atoms to increase logically. The cases often differ slightly but not too extremely because hydrogen atoms are also less stable when it comes to adsorption events. The heavy atoms (Oxygen, Carbon and Nitrogen) are therefore considered when determining the stable conformation of molecules.

**Figure 5:**
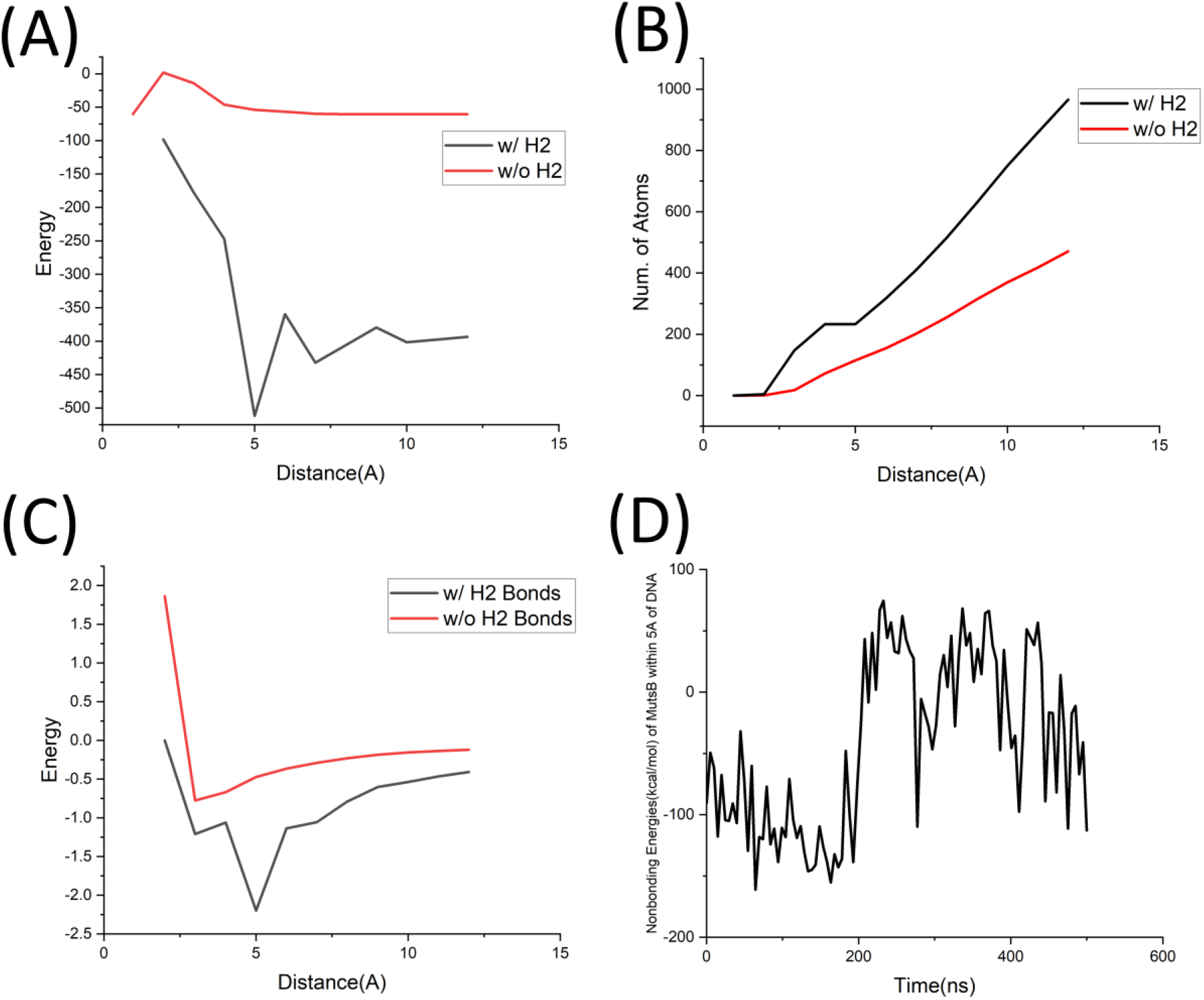
(A) Energy with and without Hydrogen Bonds (B) Number of Atoms with and without Hydrogen Bonds (C) Energy/NOA with and without Hydrogen Bonds (D) Nonbonding energy of protein within 5 angstroms of DNA

Beginning with Figure 5A, the energy is plotted at each distance from one through twelve Angstroms in the cases with and without hydrogen atoms. For the energy line including hydrogen, the energy begins slightly negative and rises back to zero. After about 5 Å, this energy then plummets almost linearly downward. The energy ends up at around -400 kCal/mol. On the other hand, the graph without hydrogen shows a steadier value around -50 Å after a similar initial spike near the 3 Å mark. Moving on to Figure 5B, this panel presents a straightforward incline for both hydrogen present and absent cases as their numbers of atoms increase when distance expands to include more atoms. By putting these two figures together, the result is Figure 5C. This plot shows the divisional relationship of energy divided by the number of atoms for both cases. The relationship between these two graphs provides the cutoff distance of ∼3 Å without hydrogen and ∼5 Å With hydrogen. This point represents the ideal absorption distance where the protein is best adsorbed onto the DNA strand as the scanning process begins. Figure 5D shows the non-bonding energy between the atoms of MutSbeta within 5 Å of the mismatched DNA and the mismatched DNA indicating the start of the scanning process at ∼200ns. As can also be observed, the non-bonding energy for atoms within 5 Å of mismatched DNA continues to drop from ∼50 kCal/mol starting ∼200ns to -100 kCal/mol by the end of the simulation run indicating that as the DNA is scanned by MutSbeta, this non-bonding energy continues to increase until the mismatched site on the DNA is reached.

As a final analysis of how DNA interacts with MutSbeta the amino acids at the last frame were collected. These are the amino acids that were adsorbed on the mismatched DNA and show what type of amino acids does the DNA prefer to interact with. As seen in table 3 the main types of amino acids that the DNA interacts the most with are polar molecules, usually polar amino acids that either have a positive or neutral charge. Specifically, lysine, arginine, asparagine, serine, cysteine, and threonine. This aligns with the fact that DNA has an overall negative charge so it is more likely to interact with molecules that are positive in nature.

**Table 3.** Adsorbed amino acids of MutSbeta on at last frame.

| Residue Name | Residue ID | Residue Type |
| --- | --- | --- |
| Lysine | 531,537,546,761,766 | Polar, positive |
| Valine | 532,752,762 | Non-polar, aliphatic |
| Arginine | 534,770 | Polar, positive |
| Asparagine | 535,536,749 | Polar, uncharged |
| Serine | 753,764,769 | Polar, uncharged |
| Cysteine | 754 | Polar, uncharged |
| Isoleucine | 755 | Non-polar, aliphatic |
| Glycine | 763 | Non-polar, aliphatic |
| Threonine | 765 | Polar, uncharged |
| Phenylalanine | 771 | Non-polar, aromatic |

## 4. Conclusions

Essential components of the relationship between MutSbeta and mismatched DNA are seen through the analysis of stability and energetics. The comparison of the RMSD’s between the MutSbeta and DNA suggests that the protein settles down as it latches onto its target DNA as it begins the repair process. The stability analysis provide solid evidence for the initial interactions between the DNA and the MutSbeta protein. After reviewing the energetic values, it is seen that the values reinforce the stability of DNA and MutSbeta when in the system as they approach each other. It also adds to the fact that the charges of the two molecules play a huge role in how they interact with each other. The exhaustive analysis determined the optimal cut-off distance of interaction between the mismatched DNA and MutSbeta to be ∼3 Å, which in-turn also demonstrated the critical number of atoms that played a crucial role in the interactions between MutSbeta and DNA. Finally, the RMSD and hydrogen bonds of the interfacial water molecules pointed out that ∼125 hydrogen bonds of the interfacial water molecules are required to stabilize the interface between MutSbeta and the mismatched DNA.

## Supplementary Materials

The following supporting information can be downloaded at: www.mdpi.com/xxx/s1, Figure S1: Clamp AP1 Secondary Structure; Figure S2: AP4 Clamp Secondary Structure; Figure S3: BP1 Clamp Secondary Structure; Figure S4: BP4 Clamp Secondary Structure; Figure S5: RMSD of DNA and MutSb of the system from 0-500ns; Figure S6: Nonbonding energy of MutSb of the system from 0-500ns; Figure S7: Electrostatics of the mismatched DNA of the system from 0-500ns; Figure S8: Electrostatics of MutSb of the system from 0-500ns.

## Author Contributions

Conceptualization, J.M. and I.M.; methodology, J.D. and J.M.; software, J.D., J.M. and I.M.; validation, J.D., J.M. and I.M.; formal analysis, J.D. and J.M.; investigation, J.D., J.M. and I.M.; resources, I.M.; data curation, J.D. and J.M.; writing—original draft preparation, J.D. and J.M.; writing— review and editing, I.M.; supervision, I.M.; project administration, I.M. All authors have read and agreed to the published version of the manuscript.

## Funding

This research received no external funding.

## Institutional Review Board Statement

Not applicable.

## Informed Consent Statement

Not applicable.

## Data Availability Statement

Modeling and Simulation software used in this study, VMD (Visual Molecular Dynamics) and NAMD (Nanoscale Molecular Dynamics), is freely available from the Theoretical and Computational Biophysics group at the NIH Center for Macromolecular Modeling and Bioinformatics at the University of Illinois at Urbana-Champaign, http://www.ks.uiuc.edu/Research/vmd/ (accessed: 11 September 2020). Atomic coordinate (protein data bank, PDB) file for the MutSbeta protein, 3THY, was obtained from the database located at the Research Collaboratory for Structural Bioinformatics (RCSB), www.rcsb.org (accessed: 15 September 2020). To create the protein structure files (PSF) and to carry out all-atom simulations, the necessary topology and force field parameter files were obtained from the Chemistry at Harvard Macromolecular Mechanics (CHARMM) database located at the MacKerell Lab at the University of Maryland, School of Pharmacy, http://mackerell.umaryland.edu/charmm_ff.shtml (accessed: 30 September 2020). A 50-51 base pair strand of DNA was modeled using an online DNA modeler tool called: The Sequence Manipulation Suite, https://www.bioinformatics.org/sms2/ (accessed: 15 October 2020) and VMD was used to create the mismatched base pair by editing the PDB file, specifically the mismatch site, the Resid84 on the mismatched DNA. The models for the three systems, mismatched DNA control, MutSbeta control and the combined system along with the parameter and configuration files are also available as part of the supporting information.

## Acknowledgments

We thank Fairfield University for providing computational resources to carry out the simulations.

## Conflicts of Interest

The authors declare no conflict of interest.

## Notes

### Competing Interest Statement

The authors have declared no competing interest.

## References

1. Zhang Y, Gong F. DNA repair. Methods. 2009 May;48(1):1–2.

2. Sclafani RA, Holzen TM. Cell Cycle Regulation of DNA Replication. Annu Rev Genet. 2007 Dec 1;41(1):237–80.

3. Chatterjee N, Walker GC. Mechanisms of DNA damage, repair, and mutagenesis. Environ and Mol Mutagen. 2017 Jun;58(5):235–63.

4. Lamers MH, Georgijevic D, Lebbink JH, Winterwerp HHK, Agianian B, De Wind N, et al. ATP Increases the Affinity between MutS ATPase Domains. Journal of Biological Chemistry. 2004 Oct;279(42):43879–85.

5. Young SJ, West SC. Coordinated roles of SLX4 and MutSβ in DNA repair and the maintenance of genome stability. Critical Reviews in Biochemistry and Molecular Biology. 2021 Mar 4;56(2):157– 77.

6. Kunkel TA, Erie DA. DNA MISMATCH REPAIR. Annu Rev Biochem. 2005 Jun 1;74(1):681–710.

7. Fishel R. Mismatch Repair. Journal of Biological Chemistry. 2015 Oct;290(44):26395–403.

8. Gupta S, Gellert M, Yang W. Mechanism of mismatch recognition revealed by human MutSβ bound to unpaired DNA loops. Nat Struct Mol Biol. 2012 Jan;19(1):72–8.

9. Fasman GD. Handbook of Biochemistry [Internet]. 0 ed. CRC Press; 2019 [cited 2023 Oct 25]. Available from: https://www.taylorfrancis.com/books/9781351080842

10. Boland CR. DNA Mismatch Repair. In: Rodriguez-Bigas MA, Cutait R, Lynch PM, Tomlinson I, Vasen HFA, editors. Hereditary Colorectal Cancer [Internet]. Boston, MA: Springer US; 2010 [cited 2023 Oct 25]. p. 67–85. Available from: http://link.springer.com/10.1007/978-1-4419-6603-2_4

11. Qiu R, Sakato M, Sacho EJ, Wilkins H, Zhang X, Modrich P, et al. MutL traps MutS at a DNA mismatch. Proc Natl Acad Sci USA. 2015 Sep;112(35):10914–9.

12. Liu Y, Prasad R, Beard WA, Kedar PS, Hou EW, Shock DD, et al. Coordination of Steps in Single-nucleotide Base Excision Repair Mediated by Apurinic/Apyrimidinic Endonuclease 1 and DNA Polymerase β. Journal of Biological Chemistry. 2007 May;282(18):13532–41.

13. Reardon JT, Sancar A. Nucleotide Excision Repair. In: Progress in Nucleic Acid Research and Molecular Biology [Internet]. Elsevier; 2005 [cited 2023 Oct 25]. p. 183–235. Available from: https://linkinghub.elsevier.com/retrieve/pii/S0079660304790042

14. Modrich P. MECHANISMS AND BIOLOGICAL EFFECTS OF MISMATCH REPAIR. Annu Rev Genet. 1991 Dec;25(1):229–53.

15. Hingorani MM. Mismatch binding, ADP–ATP exchange and intramolecular signaling during mismatch repair. DNA Repair. 2016 Feb;38:24–31.

16. Allen DJ. MutS mediates heteroduplex loop formation by a translocation mechanism. The EMBO Journal. 1997 Jul 15;16(14):4467–76.

17. Gupta S, Gellert M, Yang W. Mechanism of mismatch recognition revealed by human MutSβ bound to unpaired DNA loops. Nat Struct Mol Biol. 2012 Jan;19(1):72–8.

18. Salsbury FR. The molecular mechanism of DNA damage recognition by MutS homologs and its consequences for cell death response. Nucleic Acids Research. 2006 Apr 28;34(8):2173–85.

19. Burdova K, Mihaljevic B, Sturzenegger A, Chappidi N, Janscak P. The Mismatch-Binding Factor MutSβ Can Mediate ATR Activation in Response to DNA Double-Strand Breaks. Molecular Cell. 2015 Aug;59(4):603–14.

20. Rodrigo D, Limaj O, Janner D, Etezadi D, FJ García De Abajo, Pruneri V, et al. Mid-infrared plasmonic biosensing with graphene. Science. 2015 Jul 10;349(6244):165–8.

21. Zhang M, Liao C, Mak CH, You P, Mak CL, Yan F. Highly sensitive glucose sensors based on enzyme-modified whole-graphene solution-gated transistors. Sci Rep. 2015 Feb 6;5(1):8311.

22. Kuila T, Bose S, Khanra P, Mishra AK, Kim NH, Lee JH. Recent advances in graphene-based biosensors. Biosensors and Bioelectronics. 2011 Aug;26(12):4637–48.

23. Macwan I, Khan MDH, Aphale A, Singh S, Liu J, Hingorani M, et al. Interactions between avidin and graphene for development of a biosensing platform. Biosensors and Bioelectronics. 2017 Mar;89:326–33.

24. Humphrey W, Dalke A, Schulten K. VMD: Visual molecular dynamics. Journal of Molecular Graphics. 1996 Feb;14(1):33–8.

25. Stothard P. The Sequence Manipulation Suite: JavaScript Programs for Analyzing and Formatting Protein and DNA Sequences. BioTechniques. 2000 Jun;28(6):1102–4.

26. Jorgensen WL, Chandrasekhar J, Madura JD, Impey RW, Klein ML. Comparison of simple potential functions for simulating liquid water. The Journal of Chemical Physics. 1983 Jul 15;79(2):926–35.

27. Phillips JC, Braun R, Wang W, Gumbart J, Tajkhorshid E, Villa E, et al. Scalable molecular dynamics with NAMD. J Comput Chem. 2005 Dec;26(16):1781–802.

28. Patel R, Salamone G, Macwan I. The Role of Graphene Monolayers in Enhancing the Yield of Bacteriorhodopsin Photostates for Optical Memory Applications. Applied Sciences. 2021 Oct 18;11(20):9698.

29. Lu T, Chen Q. van der Waals potential: an important complement to molecular electrostatic potential in studying intermolecular interactions. J Mol Model. 2020 Nov;26(11):315.

30. Darden T, Perera L, Li L, Pedersen L. New tricks for modelers from the crystallography toolkit: the particle mesh Ewald algorithm and its use in nucleic acid simulations. Structure. 1999 Mar;7(3):R55–60.

31. Origin(Pro), “Version 2021b”. OriginLab Corporation, Northampton, MA, USA.

